# First evidence for an Amazonian insect migration in the butterfly *Panacea prola* (Lepidoptera: Nymphalidae)

**DOI:** 10.1101/2020.09.01.277665

**Authors:** Geoffrey Gallice, Riccardo Mattea, Allison Stoiser

## Abstract

Insect migrations rival those of vertebrates in terms of numbers of migrating individuals and even biomass, although instances of the former are comparatively poorly documented. This is especially true in the world’s tropics, which harbor the vast majority of Earth’s insect species. Understanding these mass movements is of critical and increasing importance as global climate and land use change accelerate and interact to alter the environmental cues that underlie migration, particularly in the tropics. Here, we provide the first evidence for an insect migration for the nymphalid butterfly *Panacea prola* in the Amazon, the world’s largest and most biodiverse rainforest that is experiencing a shifting climate and rapid forest loss.

## 1. INTRODUCTION

Animal migrations, defined as long-range movements of individuals to track resources that vary in space and time—breeding or overwintering grounds, for instance, or seasonally available food resources—have been documented in a wide range of species and across multiple biogeographic regions, particularly in larger-bodied species such as birds and mammals (Dingle 2014). However, despite the enormous potential importance of mass movements of smaller-bodied organisms, migrations of insect species remain comparatively poorly studied. Indeed, studies have shown that both numerically and in terms of biomass, insect migrations often far exceed those of birds and even large mammals (Holland et al. 2006). Notwithstanding a handful of well-known cases (e.g., Monarch Butterflies, Malcolm & Zalucki 1993; Painted Lady Butterfly, Pollard et al. 1998; Desert Locust, Symmons & Cressman 2001; Darner dragonflies, Russel et al. 1998), the dynamics and ecological implications of most insect migrations, and even whether most insects migrate at all, remain unknown. This lack of information is particularly acute in the world’s tropics, which harbor the vast majority of Earth’s species, especially insects. Understanding these mass movements is of increasing importance given accelerating global climate change, which is expected to strongly alter the environmental variables that serve as cues for animal migrations in the tropics, as well as the biotic interactions that further influence them (Shaw 2016).

To date and to our knowledge, not a single long-range migration has been described in the Amazon for any non-avian, terrestrial taxon, despite this being the world’s largest and most biodiverse rainforest (ter Steege et al. 2013). The large-scale environmental gradients that underly animal migrations elsewhere are especially pronounced across the Amazon, including strong east-west and north-south gradients in moisture and seasonality (Sombroek 2001). Migrations, therefore, are likely a common occurrence here, particularly among insects that are known in other regions to move large distances and in great numbers to track seasonal resources.

In this study we provide the first evidence of an Amazonian migration for *Panacea prola* Doubleday [1848], a cosmopolitan and highly abundant Neotropical butterfly species distributed from Mexico through Argentina, including throughout the Amazon.

## 2. METHODS

The study was conducted at Finca Las Piedras (FLP), a 54-ha biological research station located in Peru’s Madre de Dios department in the country’s southeastern Amazon region (12°13’32.03”S, 69°06’55.77”W; 240 m a.s.l.; Figure 1). This region lies in the extreme southwestern portion of the Amazon basin and is marked by a strong dry season that peaks annually from approximately June through September, during which fewer than 100 mm of rain generally fall per month (Fick & Hijmans 2017). The FLP property is located ca. 2 km from the Interoceanic Highway that bisects the region and is situated at the edge of the agricultural frontier that expanded upon completion of the highway in Peru between roughly 2005 and 2011. The section of the highway in the vicinity of FLP is generally bordered to the east and west by a several kilometer wide band of agricultural fields and degraded forest, after which primary forest cover is retained due to the presence of concessions for the sustainable extraction of Brazil nuts (*Bertholletia excelsa*) from natural forest. The property and surrounding area are covered mostly in mature, upland or ‘terra firme’ rainforest but active and abandoned agricultural fields and *Mauritia* palm swamps are also present.

**Figure 1.**
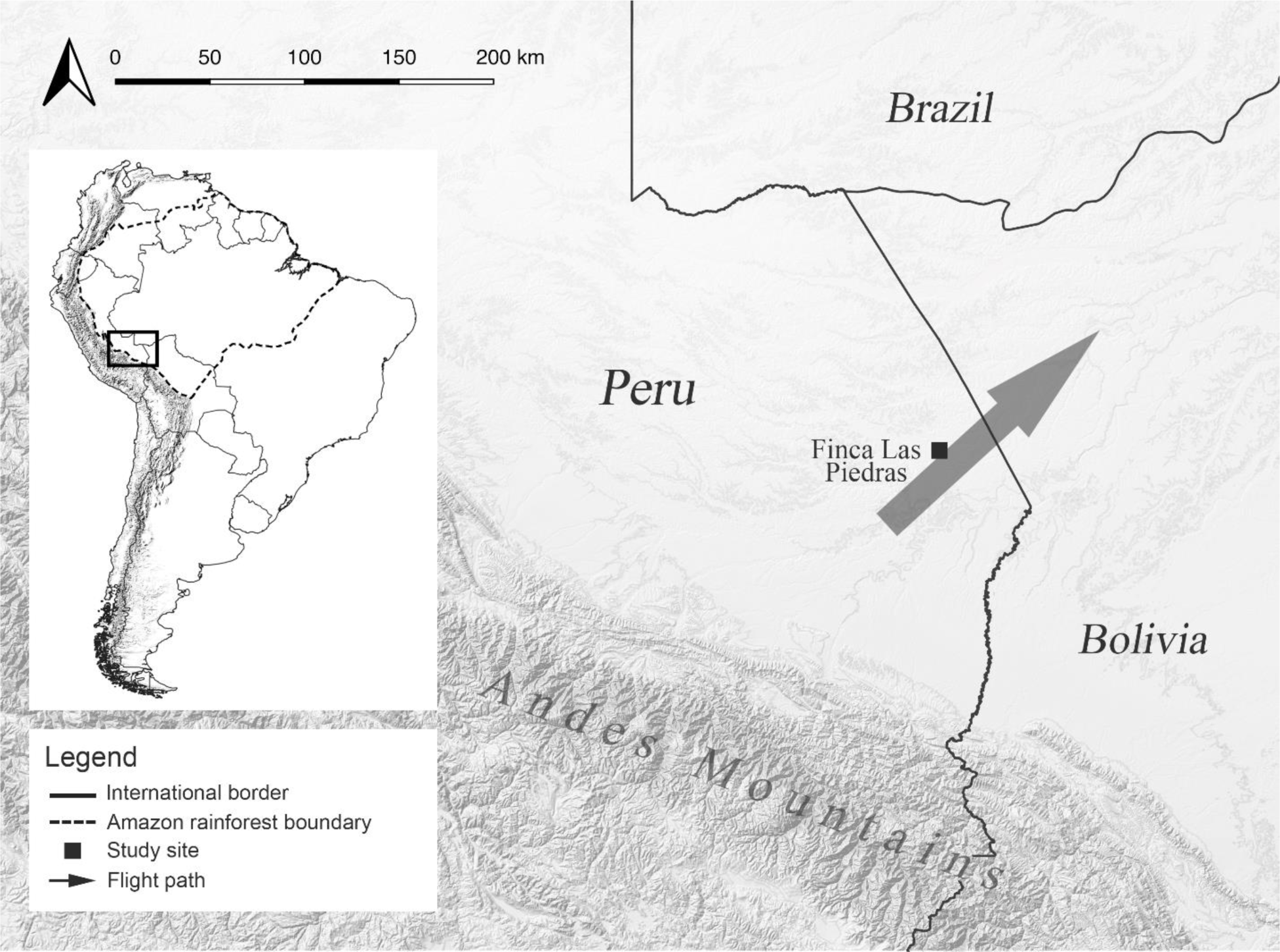
Map of the study region and migration route in southeastern Peru.

Flight behavior was recorded for adult *P. prola* at FLP during the 2019 dry season (17 August-27 October), beginning shortly after (<five days) a large number of individuals were first observed flying directionally at the field site. Sampling was conducted daily from 1200 h-1300 h for the duration of the study, which appeared to coincide with the species’ peak flight activity. An observer positioned in the NW corner of a ca. 0.5-ha clearing within a larger area of abandoned agricultural fields and regenerating secondary forest and facing approximately SE recorded individual butterflies crossing the clearing, noting the direction of flight for each individual. This position was chosen to orient the observer’s field of vision perpendicular to the butterflies’ overall flight path. Individuals were considered migratory and were counted if they displayed sustained, directional flight across the entire clearing. In addition to counts of individual butterflies, environmental data were recorded during count sessions, including cloud presence/absence and local weather (sunny, partially cloudy, cloudy, raining).

Due to excess zero counts, overdispersion, and variance much greater than the mean for the dataset, a Hurdle model was used to test the influence of recording days on the number of individuals observed. Since butterfly flight activity is known to be strongly influenced by local weather conditions (e.g., Cormont et al. 2011), a second Hurdle model was used to test the influence of cloud cover on daily counts. All data analysis was conducted using the RStudio software (RStudio Team 2015).

## 3. RESULTS

A total of 2,509 individual *P. prola* butterflies were observed displaying sustained, directional flight during 50 hours across 50 days of sampling (one hr/day), of which only 19 individuals (<0.01%) were recorded flying in a direction different from the overall flight path. Of the total count, 2,480 individuals (99%) were recorded during the first 25 days of the study. The overall flight path of migrating butterflies was approximately NE (Figure 1). Only one individual was recorded flying in a direction other than the overall flight path during the first 25 days of the study of the 19 total individuals.

The number of individuals observed during a recording day was significantly affected by recording days (Hurdle, Estimate=-0.095 *df*=49, *P*=0.001); for each recording time unit increase (recording day), on average, the probability of recording butterflies decreased by a factor of 0.095. The overall trend can be seen in Figure 2. The number of individuals observed per recording day was also significantly affected by cloud cover (Hurdle, Estimate=-1.83, *df*=49, *P*=0.027).

**Figure 2.**
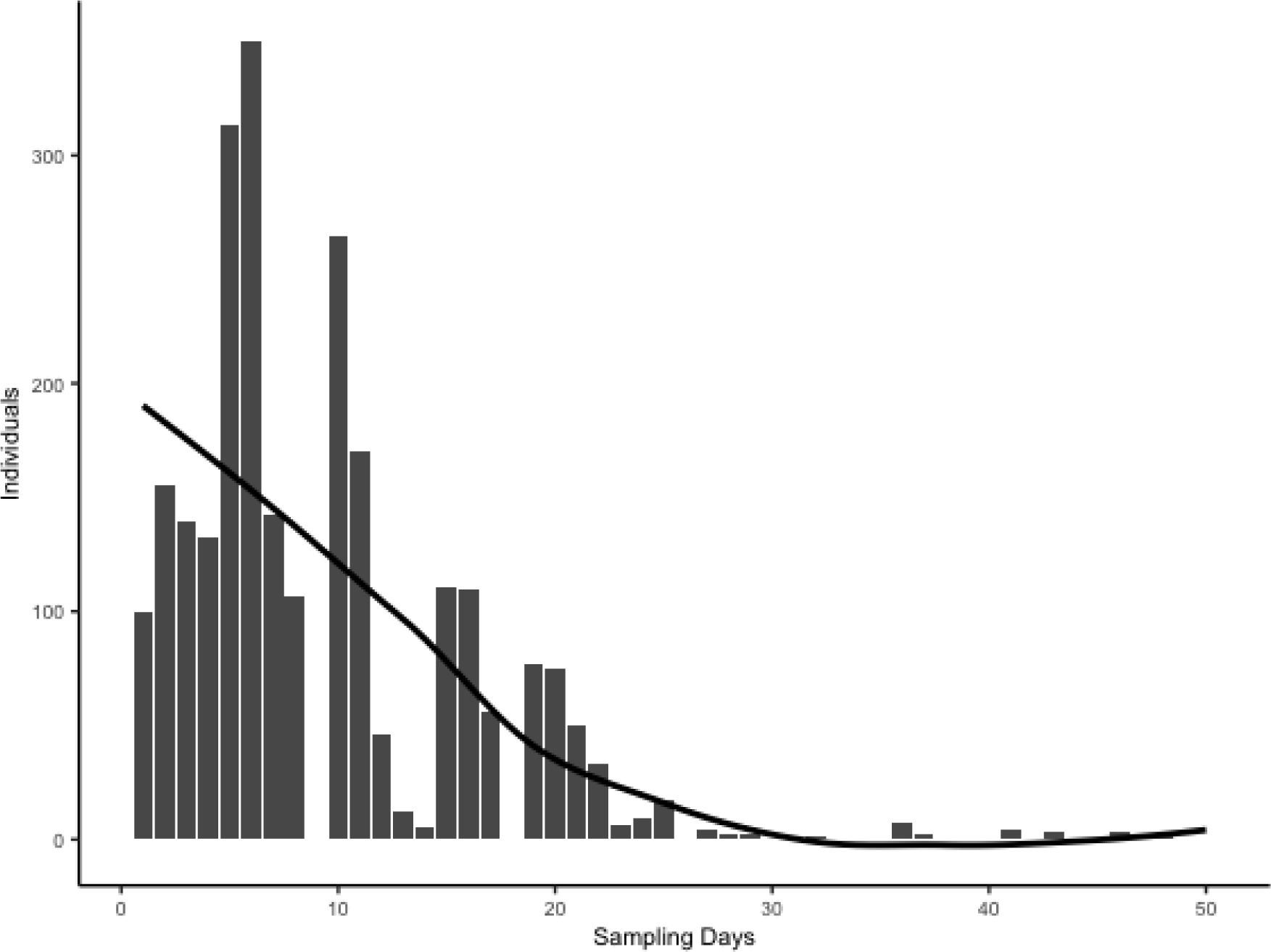
Daily counts of migrating *P. prola* individuals at the study site. Strong reductions in daily counts inconsistent with the overall trend were associated with cloud cover (Hurdle, Estimate=-1.8326, *df*=49, *P*=0.027). Recording began <5 days after the first individuals were observed flying directionally at FLP. Thick black line is the overall trend.

## 4. DISCUSSION

In this study we present the first evidence of a migration for an insect in the Amazon, the world’s largest and most biodiverse rainforest. Much of Amazonia experiences a pronounced annual dry season, particularly in the southwestern portion of the basin where this study was conducted, with potentially important implications for plant and animal population dynamics via the direct effects of seasonality on individuals’ fitness and indirect effects on species interactions. Insect herbivores such as butterflies and the plants they feed on represent a significant portion of Amazonian biomass and of total biological interactions in this ecosystem, yet how these communities and their interaction networks are influenced by seasonality is mostly unknown.

This study represents a first step in understanding these dynamics and suggests that migration is a mechanism by which at least some species cope with seasonal change in the Amazon.

Migration and diapause are common adaptations among insects to seasonality in temperate regions, where individuals of most species face a total lack of food resources and extreme environmental conditions during winter (Saulich & Musolin 2012). While seasonality and the associated declines in resource availability or quality in the tropics may not be as extreme, many tropical species do nevertheless display adaptations to seasonality. For instance, butterflies in the genus *Bicyclus* (Satyrini: Mycalesina) in the tropical African savannah switch to reproductive dormancy during times of seasonally reduced food quality (Brakefield & Reitsma 1991). Elsewhere in the tropics, migration has also been documented among insects as a means of coping with seasonal change. Srygley et al. (2010), for instance, showed for migrating *Aphrissa statira* butterflies in Panama that peak migratory behavior corresponded to dry season leaf flushing of this species’ host plant. Also in Central America, the well-documented migration of the day-flying moth *Urania fulgens* might be explained by seasonal reductions in food quality driven by an induced defensive response of host plants to herbivory (Smith 1983). Given comparable seasonal variability across Amazonia, organisms here likely face similar pressures, possibly making migration a common adaptation among Amazonian insects. However, exactly how the Amazon varies abiotically across seasons and how Amazonian organisms have adapted to this variation are more poorly understood. In the case of *P. prola*, so little is known about this species’ biology in Amazonia that a hypothesis regarding the underlying drivers of the migration documented in this study would be highly speculative at present.

Amazonia is experiencing unprecedented deforestation and forest degradation driven largely by expanding road networks and other large-scale infrastructure development (Gallice et al. 2019; Laurance et al. 2001). These impacts are compounded by global climate change which, according to some projections, may prolong the Amazonian dry season and increase forest loss through dieback and other proximate causes such as intensifying anthropogenic fire (Malhi et al. 2009). As climate change and habitat destruction interact to reduce forest cover and increase seasonal variation throughout Amazonia, the impacts on biological communities, including migration events, remain essentially completely unknown. Changes in the environmental cues that species use to decide when to migrate or to identify high quality resources might result in ecological traps if species’ behavioral changes do not keep pace with environmental change and they select habitats or other resources that do not maximize their fitness (Hale & Swearer 2016). Future research, therefore, should aim to identify other species that migrate seasonally in the Amazon, explore the underlying climatic and biotic drivers of migration in these species, and monitor changes in migration dynamics over time, to inform conservation in the rapidly changing Amazon.

## ACKNOWLEDGEMENTS

The authors would like to thank Megan Muller-Girard and Jon Pruitt for assisting with field data collection, Vicencio Oostra for providing useful feedback on the manuscript, and the Servicio Nacional Forestal y de Fauna Silvestre (National Forestry and Wildlife Service) of Peru for granting permission to conduct the field research (permit no. 187-2017-SERFOR/DGGSPFFS).

